# Neural Homophily: Similar Neural Responses Predict Friendship

**DOI:** 10.1101/092130

**Authors:** Carolyn M. Parkinson, Adam M. Kleinbaum, Thalia Wheatley

**Author notes:** Corresponding author: Carolyn M. Parkinson, Ph.D., Department of Psychology, University of California, Los Angeles, 1285 Franz Hall, Box 951563, Los Angeles, CA 90095.

## Abstract

We resemble our friends on a wide range of dimensions (e.g., age, gender), but do similarities between friends reflect deeper similarities in how we perceive, interpret, and respond to the world? To find out, we characterized the social network of a cohort of 279 students, a subset of whom participated in a functional magnetic resonance imaging (fMRI) study involving free-viewing of video stimuli. We compared fMRI response time series between corresponding brain regions across pairs of individuals and found that neural response similarity decreased with increasing distance in the social network. These effects persisted after controlling for demographic similarity. Further, it was possible to accurately classify the distance between individuals in their social network based on the similarity of their fMRI response time series across brain regions. These results suggest that we are exceptionally similar to our friends in how we perceive and react to the world around us.

## Neural Homophily: Similar Neural Responses Predict Friendship

“Birds of a feather flock together,” is an ancient truism that reflects demographic assortativity in social networks (*1*). Friends are disproportionally likely to belong to the same gender, ethnicity, and age group, and these clusters are found across diverse contexts, from traditional societies (*2*) to electronic communications (*3*) and online social networks (*4*). Demographic similarity both provides opportunities for friendship formation and influences friendship choices, given those opportunities (*5*). Here we show evidence of neural homophily above and beyond the effects of demographic similarity: when subjected to a common stimulus, friends are more similar in their neural responses than friends-of-friends who, in turn, are more similar than friends-of-friends-of-friends.

We first characterized a real-world social network among a cohort of MBA students (Fig. 1; see supplementary materials for details on social network characterization). Social distance was operationalized as geodesic distance: the smallest number of intermediary, mutual social ties required to connect two individuals in the network. Pairs of individuals in which each named the other as a friend had social distance of one. Pairs of individuals who were not friends directly but shared a mutual friend had a social distance of two, and so on.

**Figure 1.**
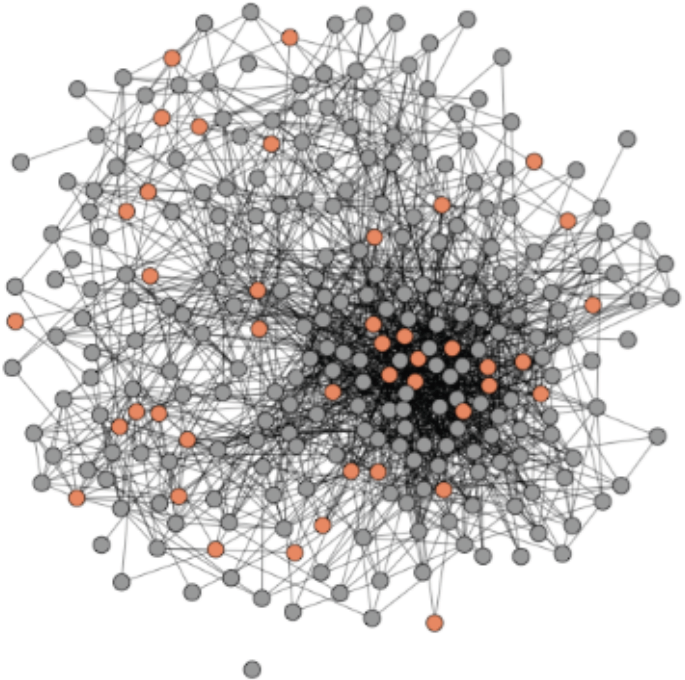
Social network. The social network of an entire cohort of first-year MBA students. Nodes indicate students; lines indicate mutually reported social ties between them. A subset of students (orange circles) participated in the fMRI study.

A subset of members of this network (*N* = 42; Fig. 1) participated in a subsequent functional magnetic resonance imaging (fMRI) study in which they viewed the same set of video clips while being scanned individually (Fig. 2b). Time series of fMRI responses measured while people view naturalistic stimuli (e.g., videos) provide an unobtrusive window into individuals’ unconstrained thought processes as they unfold (*6*). Inter-subject correlations of fMRI response time series during free viewing of naturalistic stimuli are associated with similarities in participants’ interpretations and understanding of those stimuli (*6–9*). Thus, inter-subject similarities of fMRI response time series offer insight into the similarity of individuals’ mental processing as they experience the world around them.

**Fig. 2.**
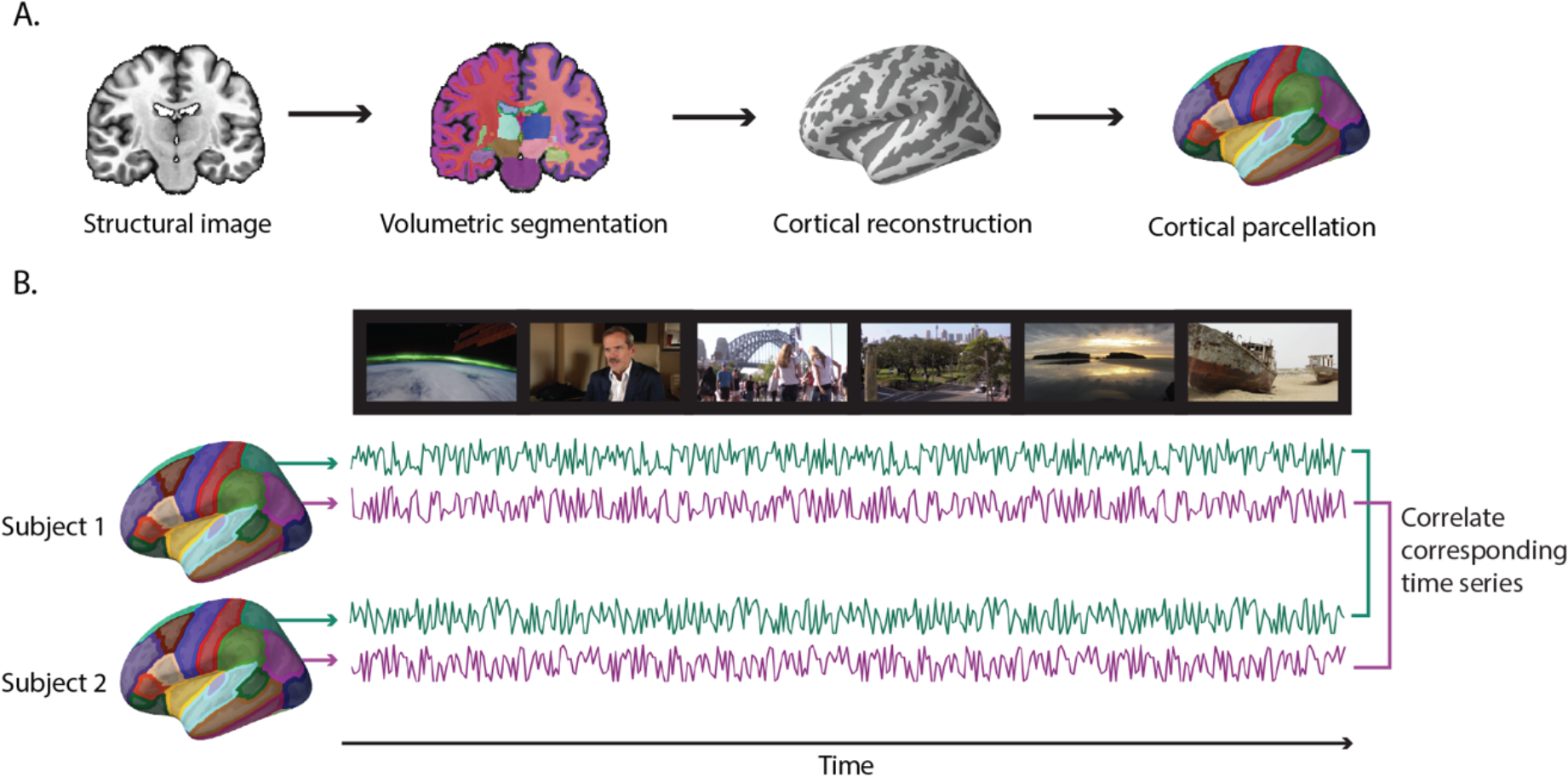
Computing inter-subject time series correlations. (A) Eighty anatomical regions of interest (ROIs) were derived for each individual using the Freesurfer image analysis suite (*10*). Segmentation of cerebral cortex, subcortical white matter, and deep gray matter volumetric structures (e.g., hippocampus, amygdala, putamen) was performed on the high-resolution scan of each individual’s brain volume. Next, a cortical surface model was reconstructed and parcellated into anatomical units. **(B)** For each individual, the average response time series within each ROI was extracted during video viewing. Next, the correlation between the time series extracted from each pair of corresponding ROIs was computed for each unique pair of participants.

The videos presented in the fMRI study covered a range of topics and genres (e.g., comedy clips, debates, documentaries; see supplementary materials). In particular, we selected videos likely to constrain participants’ thoughts and attention to the experiment (i.e., minimize mind wandering), while still promoting meaningful variability in neural response time series across participants reflective of diverging inferences, interpretations, reactions, and patterns of attentional allocation (see supplementary materials for further details).

Eighty anatomical regions of interest (ROIs) were derived for each participant using the Freesurfer image analysis suite (*10;* see supplementary materials and Fig. 2a). For each participant, in each brain region, we extracted the average fMRI response time series produced while watching the videos (Fig. 2b). Then, we computed the Pearson correlation between the time series extracted from corresponding ROIs for every possible dyad of fMRI study participants (Fig. 2b).

Of 861 possible dyads, 63 (7.32%) were characterized by a social distance of one (i.e., they were friends), 286 (33.22%) were characterized by a social distance of two (friends-of-friends), 412 (47.86%) were characterized by a social distance of three, 98 (11.38%) were characterized by a social distance of four, and two (0.23%) were characterized by a social distance of five. Given the small number of distance five dyads, data from dyads characterized by social distances of four and five were collapsed into a single category (‘4+’).

## Is social network proximity associated with similarity in neural response?

To test for a relationship between fMRI response similarity and social distance, a dyad-level regression model was used. Because the dependent variable, social distance, is ordinal, the model was estimated using ordered logistic regression. We account for the dependence structure of the dyadic data (i.e., the fact that each fMRI participant is involved in multiple dyads), which would otherwise underestimate the standard errors and increase the risk of Type 1 error (*11*), by clustering simultaneously on both members of each dyad (implemented by 3, developed by *12*). This method of accounting for dyadic dependence is comparable with approaches such as the quadratic assignment procedure or permutation testing (*13*).

Our main predictor variable of interest was neural response similarity in each student dyad. For the purpose of testing the general hypothesis that social network proximity is associated with more similar neural responses to naturalistic stimuli, neural response similarity was summarized as a single variable. Specifically, for each dyad, a weighted average of neural response similarities was computed, with the contribution of each brain region weighted by the average volume of that region in our sample of fMRI participants. (The same pattern of results obtain when weighting each ROI equally, rather than in proportion to volume.) To account for demographic differences that might impact social network structure, we also included in our model binary predictor variables indicating whether participants in each dyad were of the same or different nationalities, ethnicities and genders, as well as a variable indicating the age difference between members of each dyad. In addition, a binary variable was included indicating whether participants were the same or different in terms of handedness, given that this may be related to differences in brain functional organization (*14*).

This model revealed a significant effect of neural similarity (*β* = -0.224, *SE* = 0.106, *p* < .05) on social distance that is striking in magnitude: holding other covariates constant, compared to a dyad at the mean level of neural similarity and at any given level of social distance, a dyad one standard deviation more similar is 20% more likely to have social distance that is one unit shorter. Of the control variables also included in the model, gender (*β* = 0.383, *SE* = 0.122, *p* < .01) and nationality (*β* = 0.561, *SE* = 0.150, *p* < .001) were also related to social distance, whereas age (*β* = 0.127, *SE* = 0.136, *ns*), ethnicity (*β* = 0.094, *SE* = 0.095, *ns*) and handedness (*β* = 0.086, *SE* = 0.060, *ns*) were not. Neural similarity added significant predictive power, above and beyond observable demographic similarity (*F* = 11.06; *p* < 0.001).

Logistic regressions that combined all non-friends into a single category, regardless of social distance, yielded similar results. Neural similarity increases the likelihood of friendship dramatically: a one standard deviation increase in neural similarity increases the likelihood of friendship by 47% (*β* = 0.388, *SE* = 0.110, *p* < .001).

## Predicting friendship based on neural similarities

We also tested whether it was possible to predict friendship status based on similarity of fMRI response time series across brain regions. If so, it should be possible to build a predictive model of social distance by training an algorithm to recognize patterns of neural similarities associated with various social distance categories from a subset of dyads’ data. This model should then correctly generalize to predicting the social distances characterizing new dyads.

Eighty-element vectors of neural similarities were extracted for all 861 dyads of fMRI participants. Given that the current dataset is imbalanced across social distance categories (e.g., there are far fewer Distance 1 dyads than Distance 3 dyads), data resampling and folding procedures were used to create a series of balanced data folds such that all dyads were included in analyses (see supplementary materials for further details). Within each data fold, data were randomly partitioned into training and testing datasets. Within the training dataset for each data fold, a grid search procedure (*15*) was used to select the hyper-parameters of a support vector machine (SVM) learning algorithm that would best separate dyads according to social distance. Following hyper-parameter tuning, the classifier was trained on the entire training dataset within a given data fold to predict the social distances characterizing dyads based on corresponding patterns of inter-subject neural time course similarity. Finally, the predictive performance of this classifier was tested on data from the left-out testing dataset within the data fold, which was comprised of data from dyads to which the model had not previously been exposed. This procedure was performed within each data fold, and then cross-validated predictive performance was averaged across data folds (see supplementary materials for further details).

As shown in Figure 4, the classifier tended to predict the correct social distances for dyads in all distance categories at rates well above what would be expected based on chance alone (i.e., 25% correct). Patterns of specificity and sensitivity were consistent with good classification performance, as illustrated in the receiver-operating characteristic (ROC) curve (see Figure 4).

**Fig. 3.**
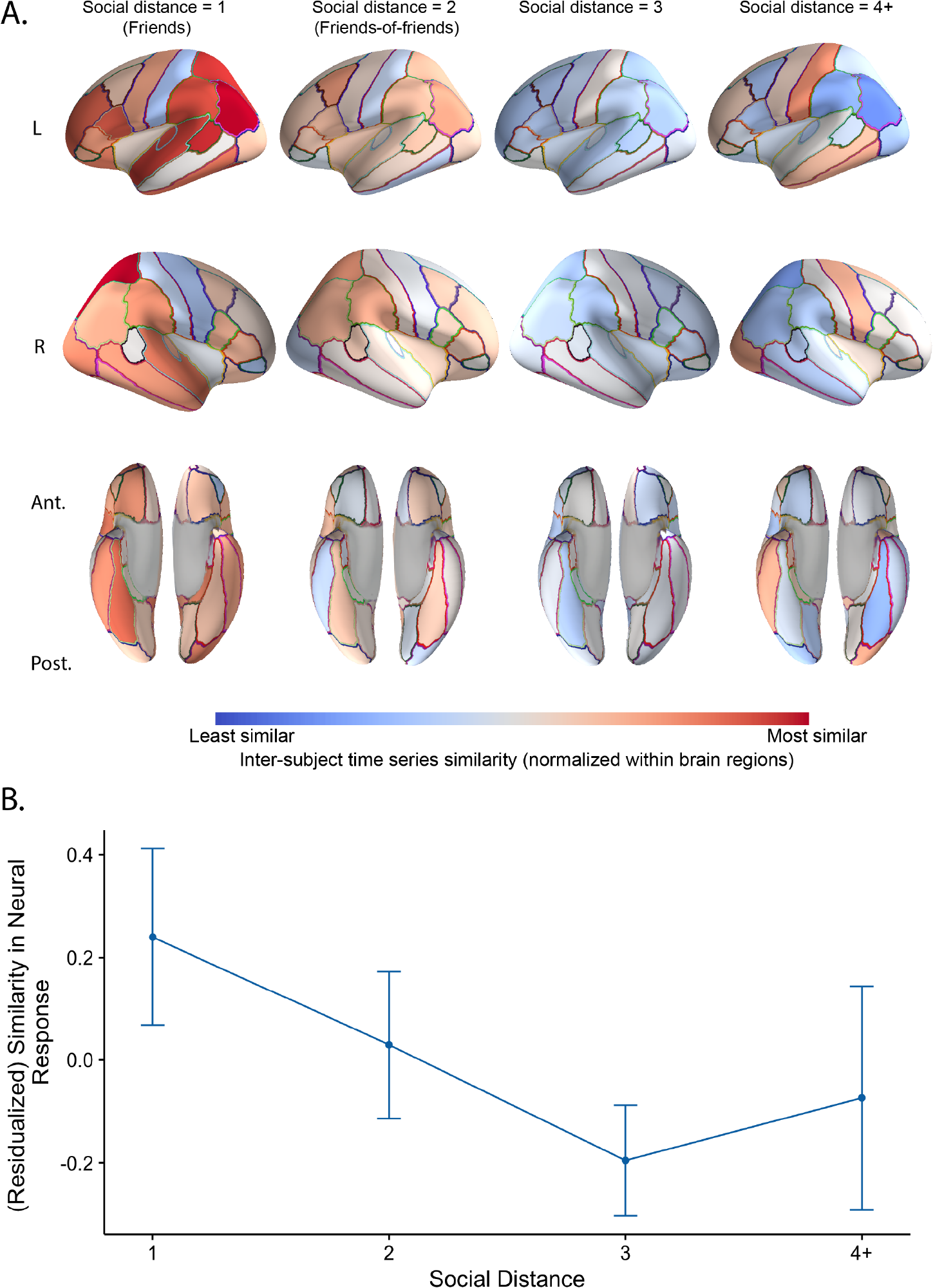
Inter-subject similarities by social distance. (A) Average dyadic fMRI response time series similarities overlaid on a cortical surface model. Average normalized inter-subject time series similarities are shown overlaid on an inflated model of the cortical surface for each social distance category. Please see supplementary materials for presentation of results that include sub-cortical gray matter structures. Ant. = anterior; Post. = posterior; L = left; R = right. **(B)** Neural response time series of dyads comprised of individuals one “degree away” from one another in the network were most similar, after residualizing out the effects of similarity in gender, nationality, ethnicity, age and handedness. Neural responses of dyads comprised of students two “degrees away” from one another were less similar (t = -2.42; p = 0.02) and those three “degrees away” were less similar still (t = -2.18; p = 0.035). Distance four dyads may be less similar (t = -1.74; p = 0.09) than distance one dyads, but were statistically indistinguishable from distance two or three. Error bars indicate 95% confidence intervals estimated using cluster-robust inference (i.e., by clustering simultaneously on both members of the dyad individually and the dyad itself).

**Fig. 4.**
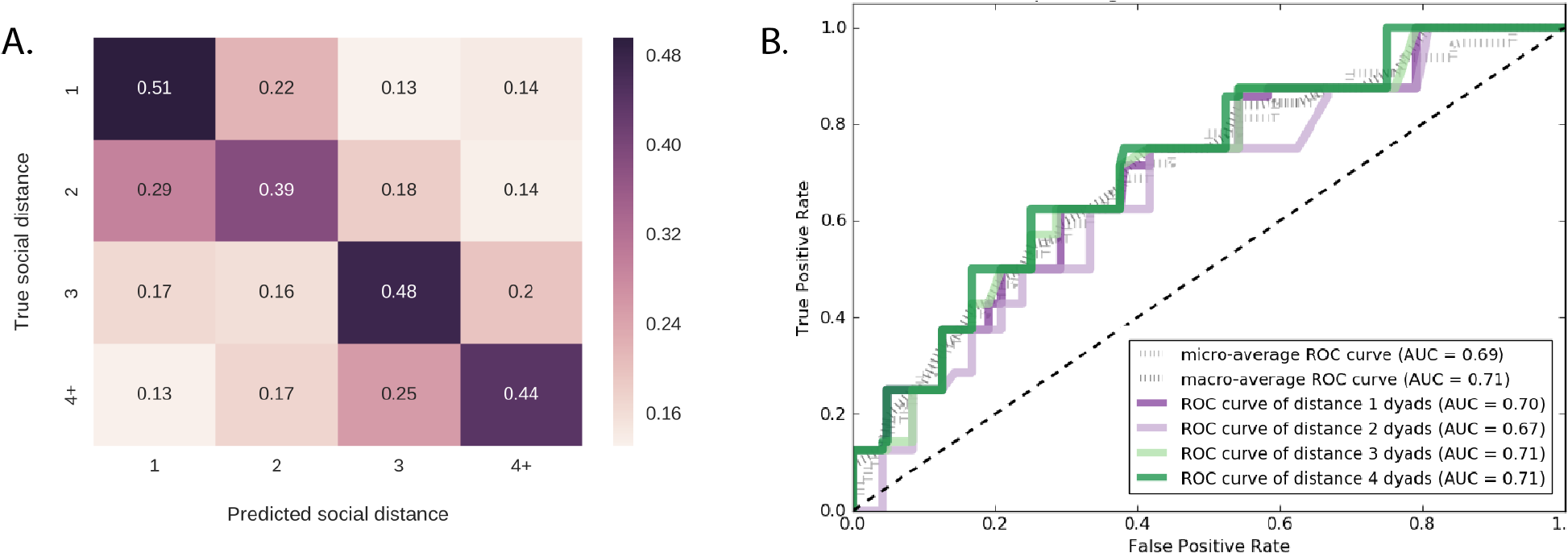
Predicting social distance based on inter-subject neural similarities. (A) Confusion matrix summarizing cross-validated prediction accuracy of four-way classifiers trained to predict the geodesic distance between members of dyads in their social network based on patterns of neural similarity, averaged across data folds (see supplementary materials for further details). Numbers and cell colors indicate how often the classifier predicted that dyads belonged to each social distance category (chance = 0.25). **(B)** Receiver-operating characteristic (ROC) curves and corresponding area under the curve (AUC) values for each social distance category. The dashed black line indicates the level of performance that would be obtained by random guessing. Points above the diagonal reflect good classification results (i.e., better than random guessing).

## Conclusion

Here we show that friends tend to respond similarly to the world around them. Neural responses during unconstrained viewing of complex, real-world stimuli were significantly more similar among friends than average. Moreover, inter-subject similarities among friends were significantly greater than inter-subject similarities among individuals at every other possible social distance from one another in the social network. In addition, predictive models trained to discern friendship status and social distance based solely on patterns of inter-subject neural response similarity were able to accurately generalize to novel data, correctly predicting the friendship statuses and social distances of new pairs of individuals based only on those dyads’ patterns of fMRI response similarities. Further, neural response similarity contributed additional explanatory power to a model predicting social distance above and beyond the contributions of demographic homophily variables, such as age, gender, nationality, and ethnicity.

Interestingly, the inverse relationship between social distance and neural similarity (as social distance increases, people have less similar brain responses) did not hold beyond 3 degrees of separation. Dyads characterized by social distances of four or more were only marginally less similar to one another than friends (i.e., distance 1 dyads) and these similarities were relatively more variable than among members of other social distance categories (Fig. 3b). There are at least two reasons why the pattern of results observed up to distance three may have ‘broken down’ at distance four. First, it is possible that individuals at distances greater than three simply do not encounter one another frequently enough to have the opportunity to become friends. Therefore, the collection of dyads characterized by a social distance of four or more may include both potentially compatible and incompatible dyads. A second, not mutually exclusive, possibility pertains to the “three degrees of influence rule” that governs diffusion in human social networks (*16*). Data from large-scale observational studies as well as lab-based experiments suggests that wide-ranging phenomena (e.g., obesity, cooperation, smoking, depression) spread only up to three degrees of geodesic distance in social networks (*16*). Although we make no claims regarding the causal mechanisms behind our findings, our results show a similar pattern.

Much previous research has shown that humans tend to associate with others who are similar to themselves in terms of a wide range of characteristics, including demographic information (e.g., age, gender, ethnicity, 1), certain personality traits and behavioral tendencies (*13,17*), and even aspects of our genotypes (*18, 19*). The current findings extend this research by demonstrating that covert mental responses to the environment, as indexed by neural processes evoked naturalistically during undirected viewing of videos, are exceptionally similar among friends.

